# US6 hijacks the peptide-loading complex by trapping transporter-chaperone dynamics

**DOI:** 10.1101/2025.06.25.661491

**Authors:** Milena Stolz, Lukas Sušac, Amin Fahim, Rieke Keller, Lisa Saggau, Filippo Mancia, Simon Trowitzsch, Robert Tampé

## Abstract

Adaptive immune responses are initiated by major histocompatibility class I (MHC I) presentation of antigenic peptides on the cell surface. This process relies on the peptide-loading complex (PLC), a dynamic transporter-multichaperone assembly in the endoplasmic reticulum (ER), to ensure high-fidelity selection, editing, and loading of peptides onto MHC I heterodimers^1^. The PLC is the primary target for viral immune evasion^2^, elicited in particular by human cytomegalovirus (HCMV)^3^, causing lifelong infections with severe risks for immunocompromised individuals. While the overall architecture of the PLC is known^4^, how its activity is jeopardized by viral immune evasins remains unclear. Here, we present the 2.59–2.88 Å cryogenic electron microscopy structure of native human PLC associated with the HCMV immune evasin US6. US6 inhibits the heterodimeric transporter associated with antigen processing (TAP1/2) by latching its transmembrane helix laterally onto TAP2 and using its central disulfide-rich domain to mimic a translocating peptide. This effectively plugs the ER-lumenal exit and locks TAP in an outward-facing open conformation with closed nucleotide-binding domains and asymmetrically occluded ATP and ADP. The structure also highlights the role of the unique N-terminal transmembrane domains of TAP as dynamic scaffolds that recruit the MHC I-specific chaperone tapasin by clamping its transmembrane helix to the core transmembrane domain of each transporter subunit. Our findings uncover the molecular mechanism of US6-mediated viral immune evasion and reveal potential targets for therapeutic modulation of antigen presentation in cancer and infectious diseases.

## MAIN

Detection and elimination of virally-infected or cancerous cells by cytotoxic T lymphocytes rely on the accurate presentation of peptides on MHC I molecules^1^, heterodimers consisting of a polymorphic MHC I heavy chain (hc) and an invariant β_2_-microglobulin (β_2_m). Central to this pathway is the peptide-loading complex (PLC), which coordinates peptide translocation into the ER with loading and editing of MHC I molecules. The PLC core consists of TAP, a heterodimeric transporter that translocates proteasomal degradation products from the cytosol into the ER lumen. Both TAP protomers are a type IV members of the conserved ATP-binding cassette (ABC) protein family, characterized by two transmembrane domains (TMDs) each consisting of six transmembrane (TM) helices and two nucleotide-binding domains (NBDs)^5^. Unique to TAP1 and TAP2 are N-terminal TM domains (TMD0s), which seem crucial for anchoring the PLC in the ER membrane and for recruiting the chaperone tapasin^6,7^, although their precise role remains elusive.

Indeed, MHC I molecules loaded with translocated peptides are edited by a network of chaperones and quality control factors, including the above-mentioned tapasin, ERp57, and calreticulin, which laterally align TAP in two editing modules^8,9^. In particular, tapasin bridges TAP with MHC I molecules. Covalently linked to ERp57^10^, it stabilizes MHC I molecules in a peptide-receptive state and facilitates the selection of high-affinity peptides through a “tug-of-war” mechanism^11,12^. Structural studies revealed that the N-terminal domains of two tapasins, composed of a seven-stranded β-barrel fused to an immunoglobulin (Ig)-like domain, recruit MHC I molecules within the ER lumen^11,13^, while their C-terminal TMDs engage TAP. Although both TMD0s can bind tapasin, TMD0 of TAP2 appears to establish a more robust interaction with the TMD of tapasin than with TAP1^14,15^. The proposed flexibility of the TAP1-tapasin interface suggests a potential division of labor within the PLC, possibly reflecting dynamic remodeling during peptide loading or release.

TAP plays a central role in adaptive immunity, and as such is often targeted by DNA viruses and downregulated in tumors^2,3^. US6 is a type-I transmembrane glycoprotein encoded by HCMV^3^, which blocks peptide translocation by TAP, without dismantling the PLC^16–21^. As a consequence, peptide-receptive MHC I molecules are retained in the ER and are consequently unable to display antigens at the cell surface to the immune system. Despite its medical relevance, for immunocompromised individuals in particular, the mechanism by which HCMV US6 inhibits TAP remains unknown. Here we investigate US6-mediated inhibition at a molecular level, in a physiological setting. This, in our opinion, offers a unique opportunity to dissect the mechanistic dependencies of the PLC and to explore how TAP’s architectural features, including the TMD0s and their interaction with tapasin, contribute to its regulation and vulnerability to viral interference.

### Structure of the US6-Arrested Human Peptide-Loading Complex

To determine the structure of a US6-arrested PLC, we utilized a monoclonal Burkitt’s lymphoma (Raji) cell line engineered for the conditional expression of US6 fused at its C terminus with a streptavidin-binding peptide (SBP)^22^ (Extended Data Fig. 1a,b). US6^SBP^ binds TAP and inhibits peptide translocation into the ER lumen (Extended Data Fig. 1a,b). After induced expression, native US6-arrested PLC was isolated from glyco-diosgenin (GDN)-solubilized Raji cell lysates via affinity chromatography and analyzed by cryo-EM (Extended Data Fig. 1-3). From a polished stack of 109,445 particles, we obtained an initial 3D cryo-EM reconstruction of the human US6-arrested PLC at an overall resolution of 3.29 Å (Extended Data Table 1 and Extended Data Fig. 3). The consensus map revealed a fully assembled PLC measuring 150 Å by 150 Å with a total height of 240 Å across the ER membrane (Fig. 1). US6-arrested PLC exhibits a tripartite architecture: the central antigen translocation module is laterally flanked by two MHC I editing modules (EM1 and EM2), each consisting of the disulfide-linked tapasin-ERp57 heterodimer, calreticulin, and an MHC I hc/β_2_m heterodimer (Fig. 1). Within these modules, tapasin, MHC I hc, and β_2_m were structurally well-resolved. Due to significant anisotropy in the consensus map, primarily caused by conformational heterogeneity in calreticulin and ERp57, we performed multiple local refinements using different focus masks to improve the resolution of individual components. These focused reconstructions yielded resolutions between 2.59 and 2.88 Å (Fig. 1, Extended Data Table 1, and Extended Data Fig. 2-4). All focused maps were aligned to the consensus map and merged into a composite map (Fig. 1 and Extended Data Fig. 2).

**Fig. 1.**
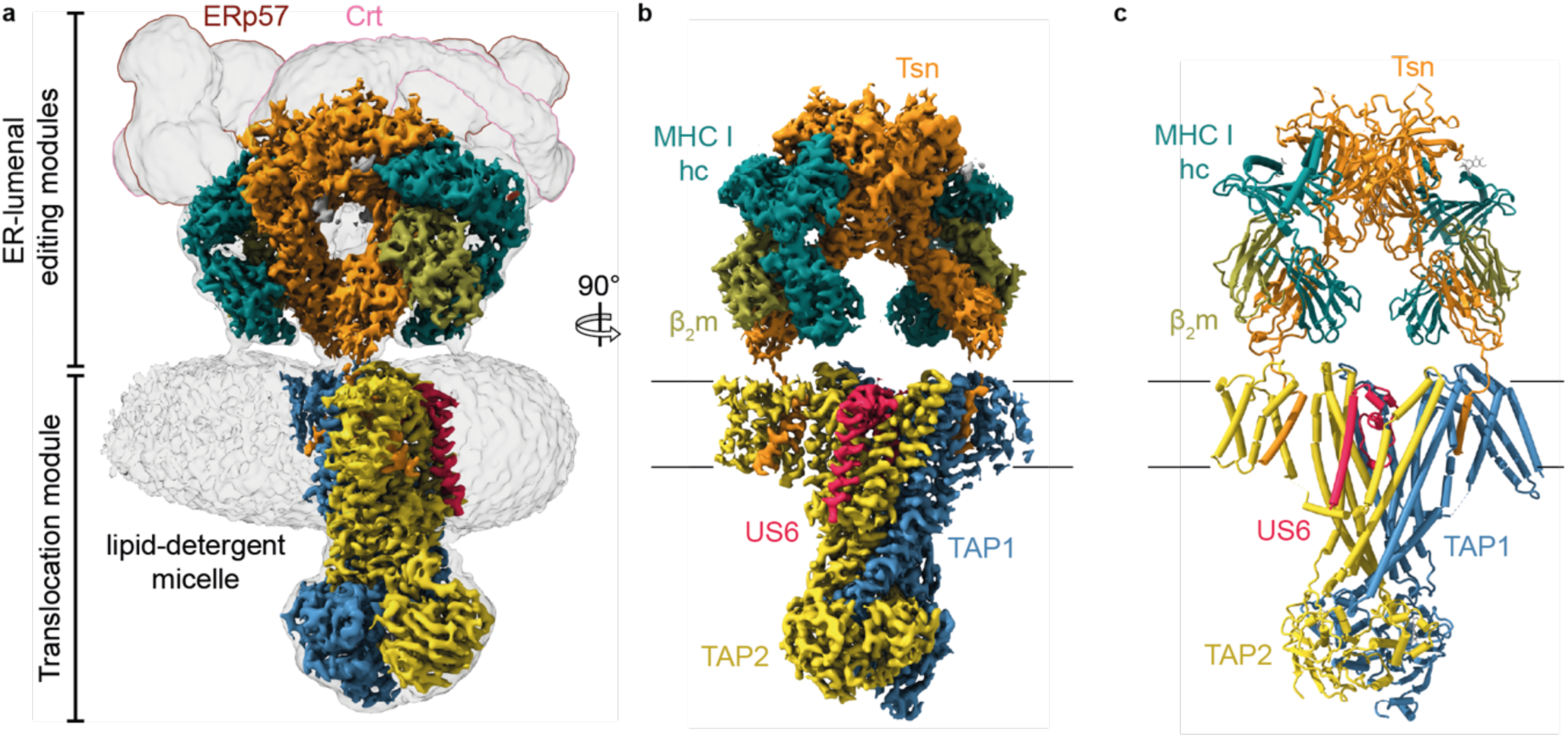
Structure of the MHC I peptide-loading complex arrested by human cytomegalovirus US6. **a**, Overview of the full peptide-loading complex (PLC), showing the lowpass-filtered consensus and the high-resolution composite cryo-EM map. **b**, Orthogonal view of the high-resolution composite map of the PLC arrested by the viral inhibitor US6. Individual subunits are labeled; subunit color coding is consistent throughout all panels. **c**, PLC structure shown as ribbon representation.

For model building, we docked structures of the editing module^13^ and the bacterial TAP-related transporter TmrAB in an asymmetric outward-facing open conformation^23^. The model of the transporter was corrected for the human TAP1 and TAP2 sequences, while the TMD0s, the TMDs of the two tapasin molecules, as well as the structure of US6 were built *de novo*. The entire assembly was refined in real space, resulting in excellent stereochemistry for the modeled entities (Fig. 1, Extended Data Fig. 3-5, and Extended Data Table 1). The final structure of US6-arrested human PLC included TAP1 (residues 17-745), TAP2 (7-683), tapasin (1-410), MHC I hc (1-274), β_2_m (1-99), and US6 (80-150). We used the primary sequence of HLA-B*15:10 to model MHC I hc in the PLC structure as immunoblotting and intensity-based absolute quantification (iBAQ) experiments showed an enrichment of the HLA-B allomorph in US6-arrested PLC^22^. The focused maps revealed the stacked back-to-back interaction of the two central tapasin molecules via electrostatic and H-bond interactions involving R28, R60, and R61 with D222, D223, and E225 (Extended Data Fig. 4e-h), similar to ICP47-inhibited PLC^4^. However, calreticulin and ERp57 could not be modeled at atomic detail given their high structural flexibility (Extended Data Fig. 2 and Movie 1).

### Topological Features of the Antigen Translocation Module

While the two editing modules of the US6-arrested PLC maintain a pseudo-symmetric configuration centered on the two tapasin molecules, US6 binding induces distinct local asymmetries. US6 anchors primarily to TAP2, altering the surrounding lipid-detergent micelle and displacing adjacent structural elements (Fig. 2). Within the lipid-detergent micelle, the core TMDs of TAP1 and TAP2 (coreTAP), along with their auxiliary TMD0s, are slightly displaced from the micelle center, likely arising from localized lipid perturbation or spatial constraints imposed by accessory proteins (Fig. 2a). Such an arrangement may reflect TAP’s physiological environment, potentially facilitating substrate access or conformational transitions in the transport cycle or optimizing the orientation of tapasin and MHC I during loading and editing. TAP adopts an outward-facing open conformation (Fig. 2b), stabilized by Mg^2+^-ADP and Mg^2+^-ATP binding at the canonical and non-canonical nucleotide-binding sites (NBSs), respectively, revealing a previously unexplored mechanism. The TMD0s exhibit a sharply angled orientation of 135° relative to the core transporter (Fig. 2c), which appears to be relevant for anchoring the editing modules and maintaining translocation fidelity.

**Fig. 2.**
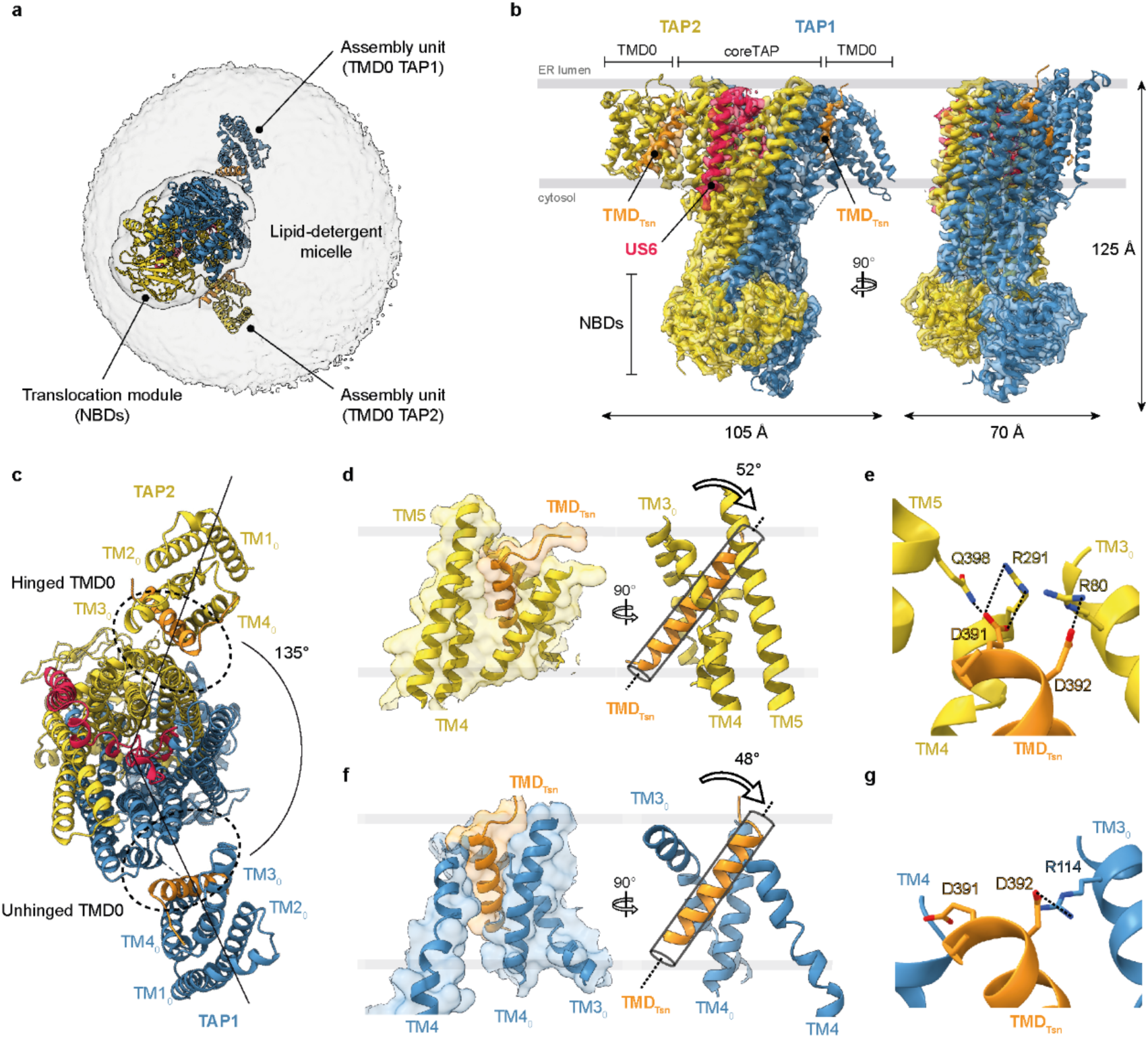
Structure of the PLC translocation module with transmembrane domains of tapasin. **a**, Cytosolic view of the peptide-loading complex showing the ribbon model of the translocation module overlaid with the lowpass-filtered consensus map. The TAP translocation module is off-center within a large lipid-detergent micelle of ∼160 Å in diameter. The lipid-detergent micelle, the two nucleotide-binding domains (NBDs) of the TAP translocation module, and the two assembly units (TMD0s) are labeled and colored as in Fig. 1. **b**, The high-resolution focused map and corresponding ribbon model of the PLC translocation module in complex with US6. The domain architecture of heterodimeric transporter TAP1 (light blue) and TAP2 (yellow) is displayed. Associated US6 (red) and the transmembrane domains of tapasin (TMD_Tsn_, orange) are indicated. Overall dimensions of the translocation module are shown. **c**, ER-lumenal view of the translocation module (as in **b**) illustrating the angled arrangement of TAP TMD0s. The interface between coreTAP and TMD0s are highlighted by dashed areas. **d**-**g**, Transmembrane domains of tapasin (TMD_Tsn_) are anchored between coreTAP and TMD0s in both TAP2 (**d**, **e**) and TAP1 (**f, g**). The tapasin transmembrane (TM) helix adopts a tilted orientation, likely to enhance the hydrophobic packing surface for interaction with TAP transmembrane segments. Two negatively charged residues at the N-terminal tip of TMD_Tsn_ engage transmembrane helices (TM) 4 and 5 of coreTAP2 and helix 3 of TAP2-TMD0. For TAP1, stabilizing interactions between tapasin and coreTAP are absent, leading to an unhinged TMD_Tsn_-TMD0 complex. Interacting residues are shown as sticks and labeled; molecular interactions are indicated with dotted lines.

An interesting feature is the robust interaction interface between TAP2, its TMD0 unit, and TMD of tapasin (TMD_Tsn_). Here, TM helix 3 and 4 of the TMD0 (TM3_0_ and TM4_0_), together with TM4 and TM5 of coreTAP, virtually enclose the TM helix of tapasin, creating an extensive interface of 1470 Å^2^ (Fig. 2c-d). These predominantly hydrophobic interactions are further stabilized by salt bridges: two acidic residues (D391 and D392) at the N-terminal tip of TMD_Tsn_ interact with basic residues R80 in TM3_0_ of TAP2’s TMD0 as well as R291 in TM4 and Q398 of coreTAP2 (Fig. 2e). These interactions stabilize a tilted orientation of the tapasin’s TM helix (52° relative to the membrane normal), anchoring this chaperone in an orientation, which appears to align with its editing function (Fig. 2d).

In contrast, the corresponding TAP1-tapasin interface (1211 Å^2^) seems weaker (Fig. 2f). It features a limited electrostatic interaction between D392 of tapasin and R114 of TAP1, suggesting a more flexible configuration (Fig. 2g). This structural asymmetry may underlie differential conformational responses during peptide translocation or MHC I peptide exchange and highlights assigned tasks between TAP1 and TAP2. Taken together, these findings support the conclusion that TAP2 plays the primary role in anchoring tapasin within the ER membrane, and thus has a greater impact on the architecture and function of the PLC than TAP1. This conclusion aligns with evolutionary differences observed in avian species, where TAP1 lacks a functional TMD0 assembly unit^24^.

### Molecular Engagement of US6 with TAP

US6 interacts with TAP through a combination of polar and nonpolar contacts. It inserts into the core TM regions of the transporter and encroaches the lumenal peptide exit gate, thereby effectively blocking antigen translocation (Fig. 3). US6 buries an extensive TAP interaction surface (3342 Å^2^), more than half the size of the TAP1-TAP2 one (6018 Å^2^). Of this, 2178 Å^2^ (65%) are accounted for the US6-TAP2 interaction. The C-terminal TM helix of US6 (residues F126-I150) nestles up against the heterodimeric transporter from the TAP2 side, lying just above and perpendicular to the TAP2 elbow helix (Fig. 3b). This arrangement is preceded by a short connecting helix (A117-H124) that wedges into a groove between TM1 and TM3 of coreTAP2 (Fig. 3b). US6 residues L80-V114 form a disulfide-stabilized domain that acts as a plug, sealing the ER-lumenal gate of the transporter (Fig. 3a,b).

**Fig. 3.**
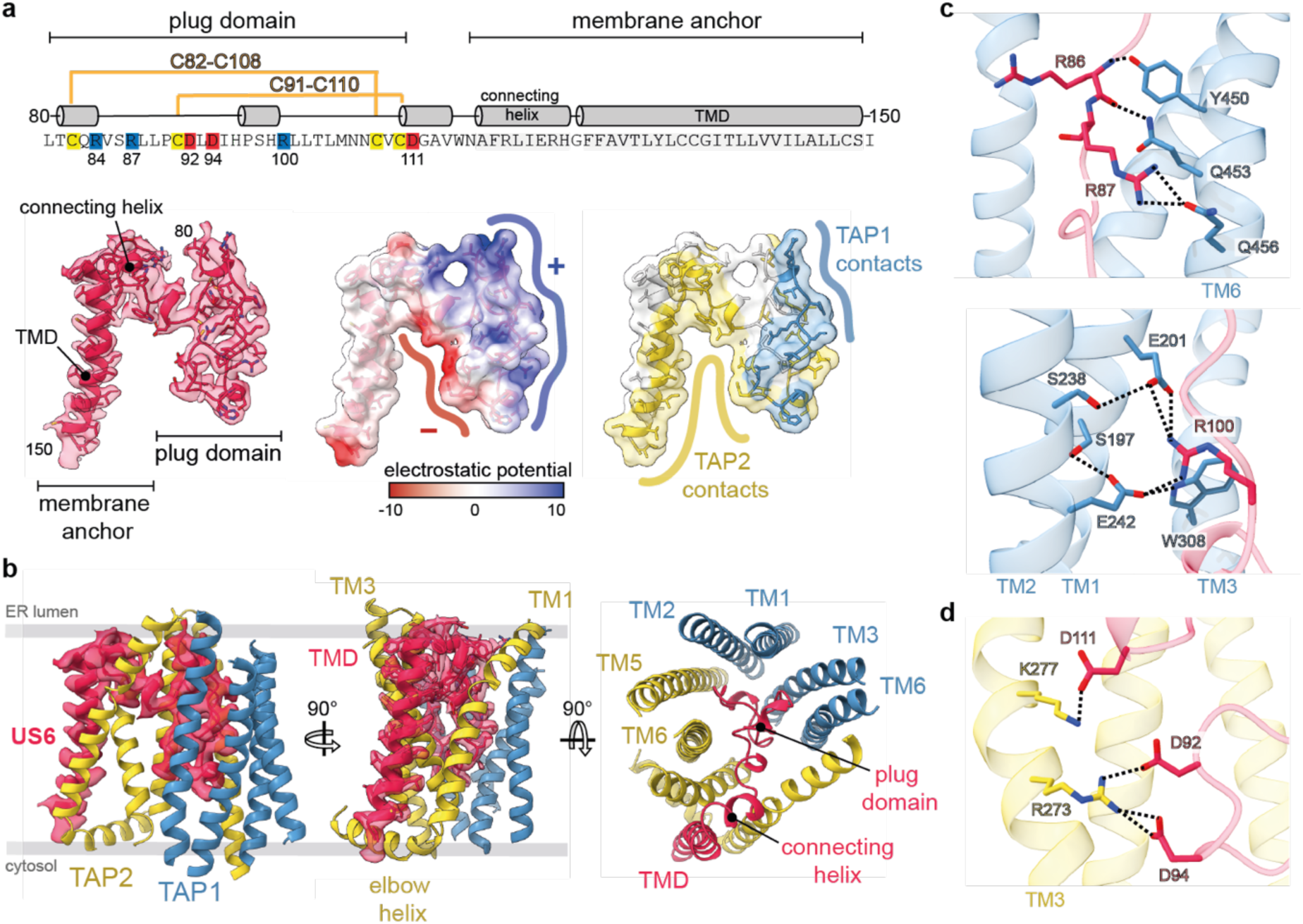
Molecular interactions of human cytomegalovirus US6 with the TAP complex. **a**, Sequence (top) and structure (bottom) of US6 interacting with coreTAP. The domain organization and secondary structure of the structurally resolved US6 region are indicated. Relevant residues are highlighted, including two intramolecular disulfide bonds (yellow) and charged amino acids (positively charged in blue, negatively charged in red). The high-resolution cryo-EM map and ribbon model of US6 (bottom left) depict the established domain organization. Charged patches in the US6 plug domain are shown as a Coulombic energy surface (bottom middle), visualizing the electrostatic potential (-10 to 10 kcal/mol). The color-coded surfaces of US6 indicate the contact sites with TAP1 (blue) and TAP2 (yellow) (bottom, right). **b**, Ribbon model of the PLC translocation module in complex with US6 showing three views of US6 interacting with adjacent coreTAP transmembrane helices. The helices are labeled accordingly. **c**, **d**, Polar interactions between the US6 plug domain and coreTAP1 (**c**) and coreTAP2 (**d**) are represented by dotted lines. Interacting residues are shown as sticks and labeled.

The connecting and TM helix of US6 engage in hydrophobic interactions with TM1 and TM3 of coreTAP2, anchoring the viral inhibitor within the membrane (Fig. 3a,b). These interactions may preclude conformational changes of the transporter, but are unlikely to effectively arrest the transport process^21^. Additional blockade appears to arise from electrostatic and steric complementarity between the charged plug domain of US6 and positively as well as negatively charged residues lining the TAP lumenal gate (Fig. 3a). Notably, the plug domain of US6 seems to neutralize key TAP residues known to coordinate the N and C termini of transported peptides^25^ (Fig. 3 and Extended Data Fig. 6). Indeed, on the TAP1 side, US6 residues R87 and R100 form polar interactions with conserved glutamate and glutamine residues (E201, E242, and Q456), which are proximal to the peptide-binding site (Fig. 3c). A mirror arrangement is observed with TAP2, where US6 residues D92, D94, and D111 form hydrogen bonds and salt bridges with R273 and K277 (Fig. 3d). This dual engagement across both protomers elegantly mimics peptide binding and locks the transporter in an outward-facing conformation.

The structural integrity of the US6 plug domain is stabilized by two disulfide bonds: C91–C110 and C82–C108 (Fig. 3a and Extended Data Fig. 6). Surprisingly, the structure reveals that the US6 plug domain does not adopt an Ig-like fold as suggested^26^. Instead, it resembles a previously unpredicted coiled-coil architecture that complements the environment of the lumenal gate in the outward-facing open conformation of TAP.

### Functional Consequences on Antigen Transport and Presentation

Our structural data support the notion that a single US6 molecule is sufficient to obstruct the peptide translocation pathway and inhibit TAP function. Although we cannot exclude the involvement of US6 multimers in additional cellular functions, our findings seem at odds with earlier models that proposed US6 multimerization as a requirement for TAP inhibition^21,27^. Comparison of the US6-bound TAP structure with the outward-facing open conformations of the homologous human transporter TAPL^28^ and the bacterial transporter TmrAB^23^ reveals a pronounced displacement of the lumenal tip of TM1 in core TAP2, likely induced by the connecting helix of US6 (Extended Data Fig. 7). As a consequence, TM5 and TM6 of coreTAP1 are slightly bent at their lumenal ends. Additionally, TM3 and TM6 of coreTAP2 adapt to the US6 plug domain, while their lumenal and cytosolic ends retain the orientation observed in the outward-facing open conformations (Extended Data Fig. 7). US6 physically occludes the translocation pore, leaving only a narrow 5 Å aperture at the lumenal gate (Fig. 4a,b). This constriction most likely prevents peptide passage and positions US6 to mimic substrate engagement, thereby locking the transporter in a non-translocating state.

**Fig. 4.**
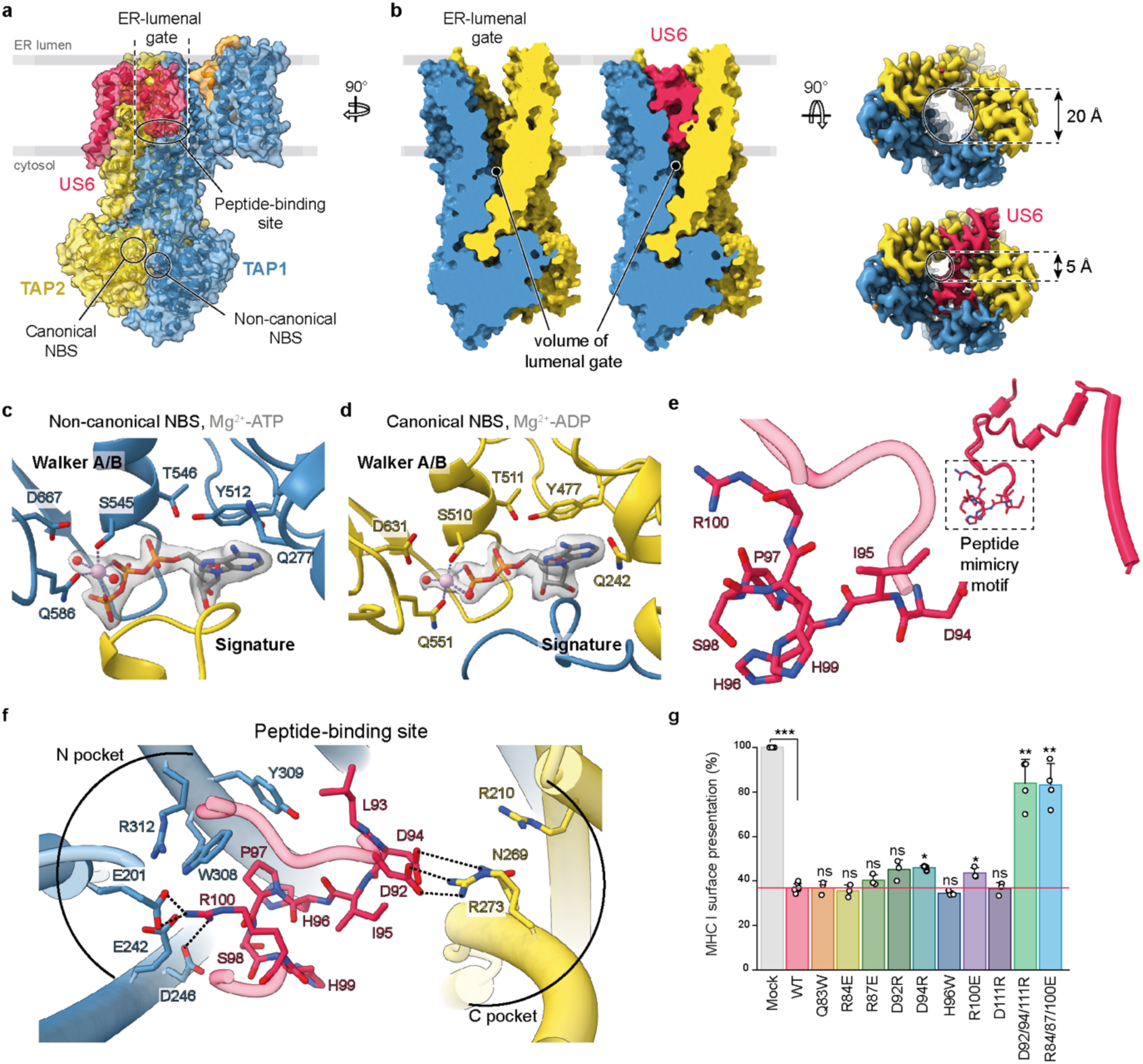
US6-mediated inhibition of TAP translocation activity. **a**, Vertical central slice through the US6-bound translocation module, highlighting structural features affected by cytomegalovirus US6-mediated inhibition. Peptide and nucleotide-binding sites (NBDs) are indicated. **b**, Outward-facing open conformation of coreTAP showing occupation of the central translocation pore by the US6 plug domain. Surface models depict 63% filling of the transporter void volume, as determined by CastpFold analysis (void volume without US6: 7495 Å^3^; with US6: 2786 Å^3^). Top views illustrate the dimensions of the lumenal gate. US6 binding does not fully seal the pore, leaving a 5 Å gap at the edge of the gate. **c, d,** Detailed views of the non-canonical (**c**) and canonical (**d**) nucleotide-binding sites (NBSs) in US6-inhibited TAP, showing bound Mg^2+^-ADP and Mg^2+^-ATP, respectively. Nucleotides are shown as sticks, Mg^2+^ ions (pink) and their coordination are indicated with dotted lines. The local EM map is displayed as a gray transparent surface. **e**, Stick and cartoon representation of the US6 peptide mimicry motif. **f**, Top view of the peptide-binding site showing US6-mediated inhibition via molecular mimicry. US6 reaches into the peptide-binding groove, occupying the binding pockets of peptide N and C terminus. Its structural polarity mimics the features of a peptide, enabling polar interactions with TAP residues critical for peptide binding. All residues are shown as sticks; interactions are marked as dotted lines. **g**, Effect of US6 mutants on MHC I surface expression. HeLa cells were transfected with mock, US6 wildtype (WT), or US6 variants. Relative MHC I surface levels (mean ± s.d., *n* = 3 forn single substitutions, *n* = 4 for triple substitutions) were normalized to mock (100%). Unpaired two-tailed *t*-tests were performed comparing mock vs. WT or individual variants vs. WT (ns, not significant; \**P* < 0.05; \*\**P* < 0.01; \*\*\**P* < 0.001).

Here, we also identified structurally well-defined nucleotides at both nucleotide-binding sites (NBSs) of the TAP heterodimer, sandwiched by the dimerized NBDs (Fig. 4 and Extended Data Fig. 7). At the canonical, catalytically active site, Mg^2+^-ADP was occluded, whereas the non-canonical inactive site coordinated Mg^2+^-ATP (Fig. 4c,d). This asymmetry suggests that the transporter is trapped in a post-hydrolysis state similarly to what was observed in TmrAB under turnover conditions^23^. The A-loop tyrosines Y512 (TAP1) and Y477 (TAP2) position the purine bases of ATP and ADP, respectively (Fig. 4c,d). The Mg^2+^ ions are coordinated by S545 and Q586 of TAP1, and by S510 of TAP2, while the coordinating sidechain oxygen of Q551 of TAP2 is displaced by 1 Å in comparison to Q586 of TAP1 (Fig. 4c,d). This displacement is linked to the hydrolysis of the β–γ phosphate bond and results in structural relaxation of the canonical NBS compared to the non-canonical NBS, as indicated by the reduced interaction interface/surface for ADP compared to ATP (429 Å^2^/546 Å^2^ and 506 Å^2^/630 Å^2^, respectively). Thus, our structural data suggest that US6 blocks return to the inward-facing conformation of TAP and nucleotide release from TAP. However, our structure cannot resolve whether TAP is captured during an active translocation cycle or in an assembly intermediate, where spontaneous ATP hydrolysis may occur at the canonical NBS. Since no exogenous nucleotides were added, the bound nucleotides are retained from the native cellular context. These findings suggest that US6 traps TAP in a post-hydrolysis state, stabilizing an asymmetric conformation with ADP and ATP bound–a key feature of its inhibitory mechanism.

The architecture of the TAP binding pocket showcases a sophisticated example of local molecular mimicry. US6 intrudes into the lumenal gate of TAP, occupying both the N- and C-terminal peptide-binding pockets^25^ (Fig. 4e,f). The electrostatic surface of the US6 plug domain closely resembles a *bona fide* antigenic peptide, allowing to engage critical substrate recognition residues of TAP (Fig. 4f). This structural mimicry seems unique among known viral evasion strategies and may represent an evolved, extremely efficient mechanism to evade immune detection while minimizing perturbation of TAP structure.

To evaluate the physiological impact of the US6-TAP interaction, we quantified MHC I surface expression in HeLa cells transfected with wild-type US6 or charge-reversal mutants targeting key TAP-interacting sites (Fig. 4g). Flow cytometry confirmed that wild-type US6 potently suppresses MHC I presentation, consistent with effective TAP inhibition (Fig. 4g and Extended Data Fig. 8). Due to the extensive interaction network between US6 and TAP, most single-residue substitutions in US6 had no significant effect compared to the wild-type protein. Only the D94R and R100E substitutions showed a modest reduction in inhibitory activity, consistent with their roles in forming salt bridges and hydrogen bonds with TAP1 and TAP2, respectively (Fig. 4f,g). Finally, the triple mutants D92/94/111R and R86/87/100E restored MHC I surface expression to near-normal levels, underscoring the essential and cooperative nature of these interactions (Fig. 4f,g).

## Discussion

Inhibition of the PLC activity by US6 unveils a previously unrecognized paradigm in viral immune evasion. By inserting into the TAP translocation pore from the ER-lumenal side, US6 effectively blocks peptide transport, disrupts MHC I loading, and silences antigen presentation at the cell surface, with an apparent minimal perturbation of the TAP structure. Our structural and functional data reveal that this effective inhibition is orchestrated through an elegant combination of electrostatic complementarity, hydrophobic membrane anchoring, and molecular mimicry of antigenic peptides. These insights highlight the extraordinary plasticity of the PLC not only in accommodating physiological substrates but also in succumbing to pathogenic inhibitors.

Surprisingly, HCMV US6 and *herpes simplex* virus ICP47 target two distinct conformations of TAP, from opposite sides^3,9,29^. US6 recognizes the outward-facing open conformation from the ER lumen whereas ICP47 wedges into the inward-facing conformation from the cytosol^3,9,29^, pinpointing TAP as a central and vulnerable checkpoint in the antigen presentation pathway (Extended Data Fig. 9). This convergence of viral strategies on a single molecular target underscores its immunological relevance and therapeutic potential.

The US6-TAP interface offers a structural blueprint for the rational design of modulators of antigen processing. Such tools could be therapeutically valuable in diseases marked by aberrant MHC I expression, including autoimmunity, viral infection, and ultimately cancer. US6 simultaneously engages both membrane-embedded and ER-lumenal surfaces of TAP, allowing for fine tuning of TAP activity by combinatorial approaches that target multiple interaction sites, similar to photo-conditional *herpes simplex* virus ICP47 (ref.^30^).

Finally, the presence of dual TMD0 assembly units has been a longstanding enigma in PLC architecture. While TAP2 TMD0 appears to serve as the primary interface with tapasin, the presence of a second TMD0 – as seen in humans – may provide advantages in scaffolding, subunit organization, or conformation regulation. Our data support a division-of-labor model, in which the TAP1-TMD0 assembly unit offers a dynamic platform for peptide editing, whereas the TAP2-TMD0 serves as the main anchor for PLC assembly, enabling iterative peptide proofreading. The sequestration of the peptide supplier TAP, the chaperone tapasin, and the peptide-recipient MHC I suggests that the PLC operates as a kinetic proofreading machinery to ensure high-fidelity antigen presentation. A more complete understanding of these coordinated mechanisms will benefit from future structural studies capturing the PLC in diverse functional states.

## METHODS

### Plasmid constructs

DNA fragments encoding US6^myc^ variants and ICP47^myc^ were generated by custom gene synthesis (TWIST Bioscience, USA) and cloned into pAMI_IRES_eGFP^11^ *via* BamHI and NotI restriction sites. Final vectors carry a gentamycin resistance gene and couple production of the viral inhibitor with an enhanced green fluorescent protein (eGFP) via an interjacent internal ribosomal entry site 2 (IRES2). All vectors were confirmed by Sanger sequencing (Microsynth).

### Cell lines

Wild-type HeLa cells (ATCC, CCL-2) were cultured in Dulbecco’s Modified Eagle Medium (DMEM, Gibco), supplemented with 10% (v/v) fetal calf serum (FCS, Gibco), in a humified atmosphere at 37 °C and 5% CO_2_. At ∼70% confluency, cells were detached using pre-warmed 0.05% (w/v) trypsin-EDTA (Gibco) for 4 min at 37 °C, followed by centrifugation at 300 x *g* for 3 min. Cell pellets were washed with Dulbecco’s phosphate-buffered saline (DPBS, Gibco) at pH 7.4 and seeded in new culture dishes pre-aliquoted with culture media.

Burkitt’s lymphoma (Raji) cells (ATCC, CCL-86) were cultured in RPMI 1640 (Gibco), supplemented with 10% (v/v) FCS, 3 mM HEPES-NaOH pH 7.5 (Gibco) and 100 U/ml Penicillin-Streptomycin (PenStrep, Gibco), under humidified conditions (37 °C, 5% CO_2_). For large-scale cultivation, Raji cell culture was adapted to two-liter Erlenmeyer flasks, incubated at 37 °C with 5% CO_2_, and shaken at 100 rpm. Cell suspensions were split at a 1:2 ratio every two to three days.

### Generation of stable monoclonal Raji US6^SBP^ cell lines

Raji cells expressing US6^SBP^ were obtained via lentiviral transduction^22^. The resulting polyclonal population was induced with 5 µg ml^-1^ doxycycline and harvested 24h post-induction. Following two washes with FACS buffer (1x DPBS, 10% (v/v) FCS), cells were subjected to single-cell sorting using a Sony MA900 cell sorter operated in 3-drops single-cell sorting mode. mCherry-positive cells were gated and sorted into 96-well plates pre-filled with 300 µl RPMI media supplemented with 20% (v/v) FCS and 100 U/ml PenStrep. Single-cell clones were expanded over 2–3 weeks and subsequently transferred to 6-well plates containing 1 ml fresh culture media. Monoclonal cell lines exhibiting optimal US6 expression were adapted to large-scale suspension culture.

### Transfection of HeLa cells

5 x 10⁵ HeLa cells per well were seeded in 6-well culture plates with 2 ml of DMEM cultivation media 18 h prior to transfection. Upon reaching ∼80% confluency, the medium was removed, and cells were washed with 1x DPBS before being supplemented with 1 ml fresh DMEM. For transfection, a mix containing 1.5 µg plasmid DNA and 4.5 µl X-tremeGENE HP DNA transfection reagent (Roche) in 100 µl Opti-MEM (Gibco) was prepared and incubated at room temperature for 15 min before being added to each well. After 5 h, 2 ml of fresh DMEM was added to each well to recover transfected cells. Cells were harvested 72 h post-transfection by incubation with trypsin-EDTA for 4 min at 37 °C, followed by centrifugation at 300 x *g* for 3 min.

### Protein production

Upon reaching a density of 1 x 10^6^ Raji cells ml^-1^, small-scale protein production (up to 5 ml) was induced by directly adding 5 µg ml^-1^ doxycycline for 24–48 h. For large-scale expression, Raji cells were cultured in 800–1000 ml volumes in two-liter Erlenmeyer flasks and grown to a density of 1 x 10^6^ cells ml^-1^. One hour prior to induction, suspension cultures were incubated without agitation at 37 °C and 5% CO_2_ to allow cell sedimentation. Subsequently, 20% of the culture supernatant was replaced with fresh, pre-warmed medium, and US6^SBP^ production was induced by adding 5 µg ml^-1^ doxycycline. Cells were harvested 48 h post-induction by centrifugation at 1000 x *g*. Cell pellets were snap-frozen in liquid nitrogen and stored at -80 °C until further use.

### Biochemical analyses

For SDS-PAGE and immunoblot analysis, 11% Laemmli gels (for purified PLC) or 11% Tris-Tricine gels (for HeLa cell lysates) were used. Proteins were directly visualized using InstantBlue^TM^ (Expedeon) or transferred onto nitrocellulose or PVDF membranes (Cytiva) for antibody-based detection. The presence of individual PLC components was assessed using the primary antibodies anti-TAP1 (mAb148.3, hybridoma supernatant, 1:10), anti-TAP2 (mAb435.3, hybridoma supernatant, 1:10), anti-HLA-A/B/C (HC10, hybridoma supernatant, 1:10), anti-β_2_m (Novo Antibodies, HPA006361, 1:1000), anti-tapasin (Abcam, ab13518, 1:1000), anti-ERp57 (Abcam, ab10287, 1:2000), anti-calreticulin (Sigma, C4606, 1:2000), and anti-SBP (Santa Cruz, sc-101595, 1:100) for detection of US6^SBP^. Production of US6 variants in HeLa cells was confirmed by anti-myc immunodetection (Sigma, 05724, 1:2000), with anti-β-actin staining (Sigma, A2228, 1:2000) serving as a loading control. The integrity of PLC::GDN complexes was verified by size exclusion chromatography on an Äkta Pure Micro HPLC system (Cytiva) equipped with a Superose 6 3.2/300 column (Cytiva), using SEC buffer (20 mM HEPES-NaOH pH 7.5, 150 mM NaCl, 0.003% (w/v) GDN) at 4 °C.

### Flow cytometry

Optimal induction of US6^SBP^ expression in stably transduced Raji cells was monitored by flow cytometry via mCherry fluorescence. For cytometry analysis, 1 x 10^6^ Raji cells were harvested 48 h post-induction and washed twice with FACS buffer (1x DPBS containing 2% (v/v) FCS) at 300 x *g* for 5 min. Following the transfection of HeLa cells, 2 x 10⁵ cells per sample were collected and washed with FACS buffer. To assess MHC I surface expression, cells were resuspended in 50 µl FACS buffer and incubated for 20 min at 4 °C with 10% (v/v) human FcR blocking reagent (Miltenyi Biotec) and Alexa Fluor 647-conjugated anti-human HLA-A/B/C antibody (clone W6/32; BioLegend; final concentration 14 nM). Cells were extensively washed and resuspended in 250 µl FACS buffer prior to cytometry analysis.

MHC I surface expression and US6^SBP^ induction in Raji cells were analyzed on a SH800S Cell Sorter (Sony) and NovoCyte Flow Cytometer (Agilent), respectively, using mCherry as a reporter. Transfection efficiency and MHC I surface expression in HeLa cells were assessed on a FACSCelesta flow cytometer (BD), using eGFP as a transfection marker. Data were processed using FlowJo v10.10.0 (TreeStar). Geometric mean fluorescence intensities (MFI) were calculated, normalized to respective controls, and compared between conditions. The gating strategy is described in Extended Data Fig. 8.

### Purification of PLC arrested by US6

Cell pellets were thawed and resuspended in lysis buffer (20 mM HEPES-NaOH pH 6.5, 150 mM NaCl, 10 mM MgCl_2_), and cOmplete^TM^ EDTA-free protease cocktail (Roche). Cells were lysed in the presence of 2% (w/v) glyco-diosgenin (GDN, Anatrace) using a PTFE tissue grinder (Cole-Parmer), followed by incubation for 2 h at 4 °C under constant agitation. Insoluble material was removed by ultracentrifugation at 100,000 x *g* for 60 min at 4 °C. The clarified solubilizate was incubated with Streptavidin High-Capacity Agarose (Pierce) for 1 h at 4 °C and extensively washed. US6-arrested PLC was eluted in buffer containing 20 mM HEPES-NaOH pH 6.5, 150 mM NaCl, 0.05% (w/v) GDN, and 2.5 mM biotin. The eluate was concentrated using Amicon Ultra-15 centrifugal filters with a 100 kDa molecular weight cut-off (Merck).

### Cryo-EM sample preparation and data acquisition

For cryo-EM grid preparation, 3 µl of purified US6^SBP^-PLC complex (3.5 mg ml^-1^, 5.6 µM) was applied to a glow-discharged holey gold grid (UltrAuFoil R0.6/1 Au 300-mesh, Quantifoil) using an easiGlow^TM^ discharge unit (PELCO). Grids were blotted for 5 s with a blot force of 5 at 4 °C and 100% humidity in a Vitrobot Mark IV (Thermo Fisher Scientific). Samples were vitrified by plunge-freezing into liquid ethane. A total of 25,454 micrographs were automatically collected on a Titan Krios transmission electron microscope (FEI) operated at 300-kV and equipped with a K3 direct electron detector (Gatan), using Leginon for automated data collection. Micrographs were recorded at a nominal magnification of 105,000x, corresponding to a calibrated pixel size of 0.83 Å. Movies were acquired with a total electron dose of 58 e^-^/Å^2^ over a defocus range from -0.5 to -1.5 µm.

### Cryo-EM data analysis

Cryo-EM data processing of 25,454 micrographs was performed using cryoSPARC v4.6.2 (ref.^31^), as summarized in Extended Data Fig. 2 and 3. Patch-motion correction and patch-based CTF estimation were carried out using the implemented jobs in cryoSPARC. Initial, exploratory processing involved blob picking and extraction of particles in a binned box size of 128 pixels. After 2D classification, prominent US6-PLC classes were re-extracted at a full box size (512 pixels) and used as templates for template-based particle picking. Based on particle count, selective high-quality micrographs were used to train the Topaz neural network. In total, 726,723 particles were picked with Topaz and subjected to *ab-initio* heterogeneous reconstruction using five classes. The class presenting two editing modules (269,213 particles) was used for initial non-uniform refinement to generate a universal starting structure for further processing.

For the consensus refinement, a three-class heterogeneous refinement was performed to isolate a homogeneous subset of 109,445 particles representing both, the two editing modules and one translocation module of the PLC. The consensus map was obtained via non-uniform refinement without alignments. Multiple local refinements using different focus masks were carried out to enhance regional density features. Focus maps were aligned to the consensus map and merged using voxel-wise maximum intensity projection in USFC Chimera^32^.

### Model building

Structures of the editing module subunits tapasin, β_2_-microglobulin, and HLA-B*15:10 (PDB ID: 7QPD) were initially docked into the cryo-EM map using UCSF Chimera and subsequently rebuilt against the corresponding focus maps in Coot^33^. Glycosylation sites were built using the Glyco-Builder tool implemented in Coot. For modeling the TAP heterodimer, the outward-facing open conformation of TmrAB (PDB ID: 6RAH^23^) was automatically docked into the TAP-specific focus map, mutated to the TAP1/2 sequence, and manually adjusted in Coot. The transmembrane helices of tapasin from both editing modules, as well as US6, were modelled *de novo* and adjusted in Coot. Final models were subjected to automated real-space refinement in Phenix^34^ using the minimization_global strategy, followed by manual rotamer corrections in Coot. Final refinement statistics were generated in Phenix and are summarized in Extended Data Table 1. Macromolecular interfaces were determined by PISA analysis^35^. Void volumes of the TAP lumenal gate were investigated by CASTpFold analysis^36^.

### Statistical analyses

Statistical analysis was performed using Excel Data Analysis tool. Flow cytometric detection of MHC I was analyzed by unpaired, two tailed *t* tests. Statistical tests and *P* values are reported in the figure legends.

## DATA AVAILABILITY

The cryo-EM density maps of the US6-arrested PLC have been deposited in the Electron Microscopy Data Bank under accession numbers: EMD-53326, EMD-53330, EMD-53331, EMD-53332, and EMD-53334. Atomic coordinates for the atomic models have been deposited in the Protein Data Bank (http://www.rcsb.org) under accession number PDB ID 9RCV. All other data are available from the corresponding author upon reasonable request. Source data are provided with this paper.

## ACKNOWLEDGMENTS

Cryo-EM sample preparations were screened at the Glacios cryo-transmission electron microscope (Thermo Fisher Scientific) of the Cryo-EM Infrastructure within the Collaborative Research Center (CRC 1507/Z02). Cryo-EM data were finally collected at the Columbia University Cryo-Electron Microscopy Center. This work was generously supported by the Schaefer Research Scholars Program from Columbia University (to R.T.). This work was also supported by the European Research Council (ERC Advanced Grant 101141396 to R.T.), the German Research Foundation (DFG Grant TA157/12-1 to R.T.), and the Collaborative Research Center CRC 1507 (P18 and Cryo-EM Infrastructure Z02 to R.T.). We thank Oliver Clarke (Departments of Anesthesiology, and Physiology & Cellular Biophysics, Columbia University) and Yudhajeet Basak (Institute of Biochemistry, Goethe University) for sharing their helpful advice in data processing and model building. We thank the members of the Filippo Mancia lab, including Brian Kloss, Jonathan Kim, and Allen Zinkle, members of the Oliver Clarke lab, including Francesca Vallese and Kookjoo Kim, and members of the cryo-EM facility, including Robert Grassucci, Zhening Zhang, and Yen-Hong Kao. We are grateful to all members of the Institute of Biochemistry, Goethe University Frankfurt for discussion and Inga Nold and Andrea Pott for editing of the manuscript.

## AUTHOR CONTRIBUTIONS

Cryo-EM data processing: MS, LSu; Methodology: MS, LSu, AF, RK, LSa, ST, RT; Investigation: MS, AF, RK, LSa, ST, RT; Visualization: MS, RT; Writing – original draft: MS, ST, RT; Writing – review & editing: MS, LSu, AF, F.M., ST, RT; Conceptualization: ST, RT; Funding acquisition: RT; Supervision: RT.

