## Supplemental Figure 1-9 and Table 1 for "US6 hijacks the peptide-loading complex by trapping transporter-chaperone dynamics"

### Extended Data Figures

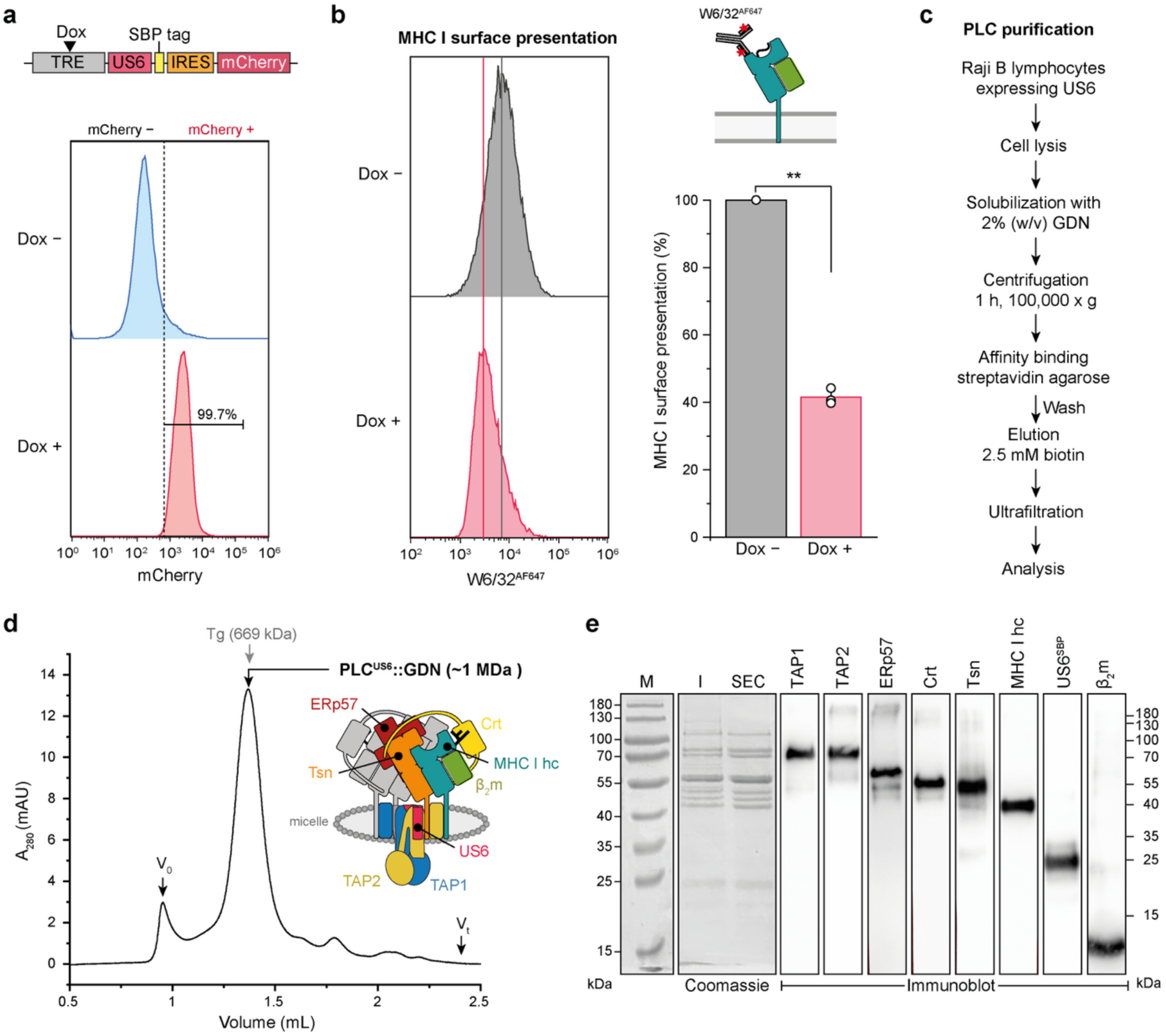


**Extended Data Fig. 1 | Induced US6 expression and purification of the native, fully assembled peptide-loading complex. a**, A monoclonal Raji cell line was engineered to conditionally express US6, which was C-terminally fused to a streptavidin-binding peptide (SBP)[22]. The lentiviral construct features a tetracycline-responsive element (TRE) driving bicistronic expression of US6$^{SBP}$ and the auto-fluorescent reporter protein mCherry via an internal ribosome entry site (IRES). US6$^{SBP}$ expression was induced with doxycycline (± Dox) and monitored by flow cytometry. **b**, Doxycycline-induced expression of US6 resulted in downregulation of MHC I surface expression, assessed by flow cytometry using an Alexa Fluor 647-conjugated anti-HLA-A/B/C antibody (W6/32). **c**, Schematic of the purification workflow. **d**, The US6-arrested PLC was isolated from glyco-diosgenin (GDN)-solubilized Raji cell lysates via affinity chromatography. Its monodispersity was evaluated by size exclusion chromatography. **e**, The composition of the US6-inhibited PLC was analyzed by SDS-PAGE (11% Tris-Tricine gels, reducing conditions) and subsequent immunoblotting.

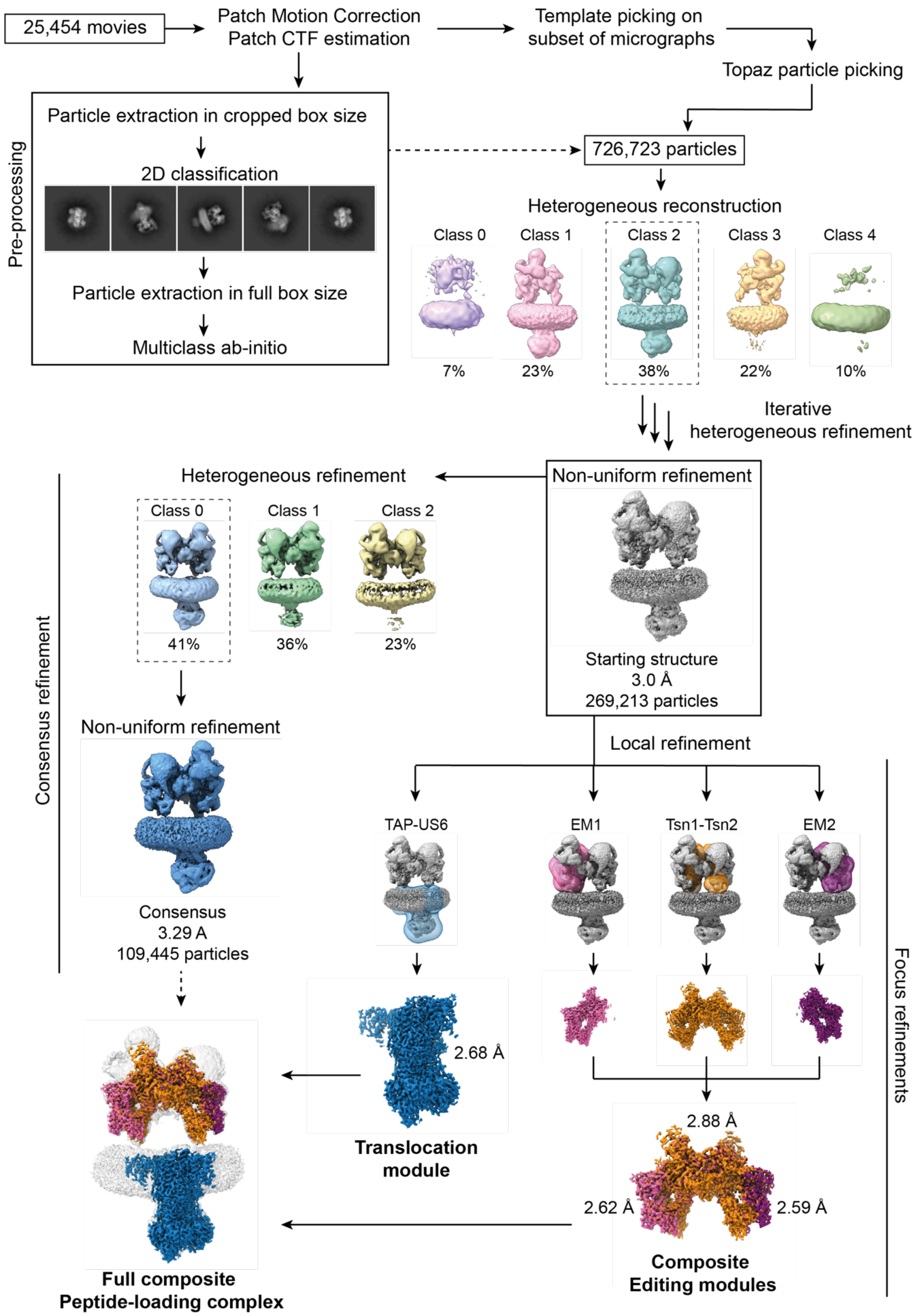


**Extended Data Fig. 2 | Cryo-EM processing and analysis workflow of the peptide-loading complex.** Flowchart illustrating the cryo-EM processing steps. All processing was performed using CryoSPARC v.4.6.2. Focus maps of Tsn-MHC I hc-$\beta_2$m complexes from editing module 1 and 2 are named EM1 and EM2, respectively.

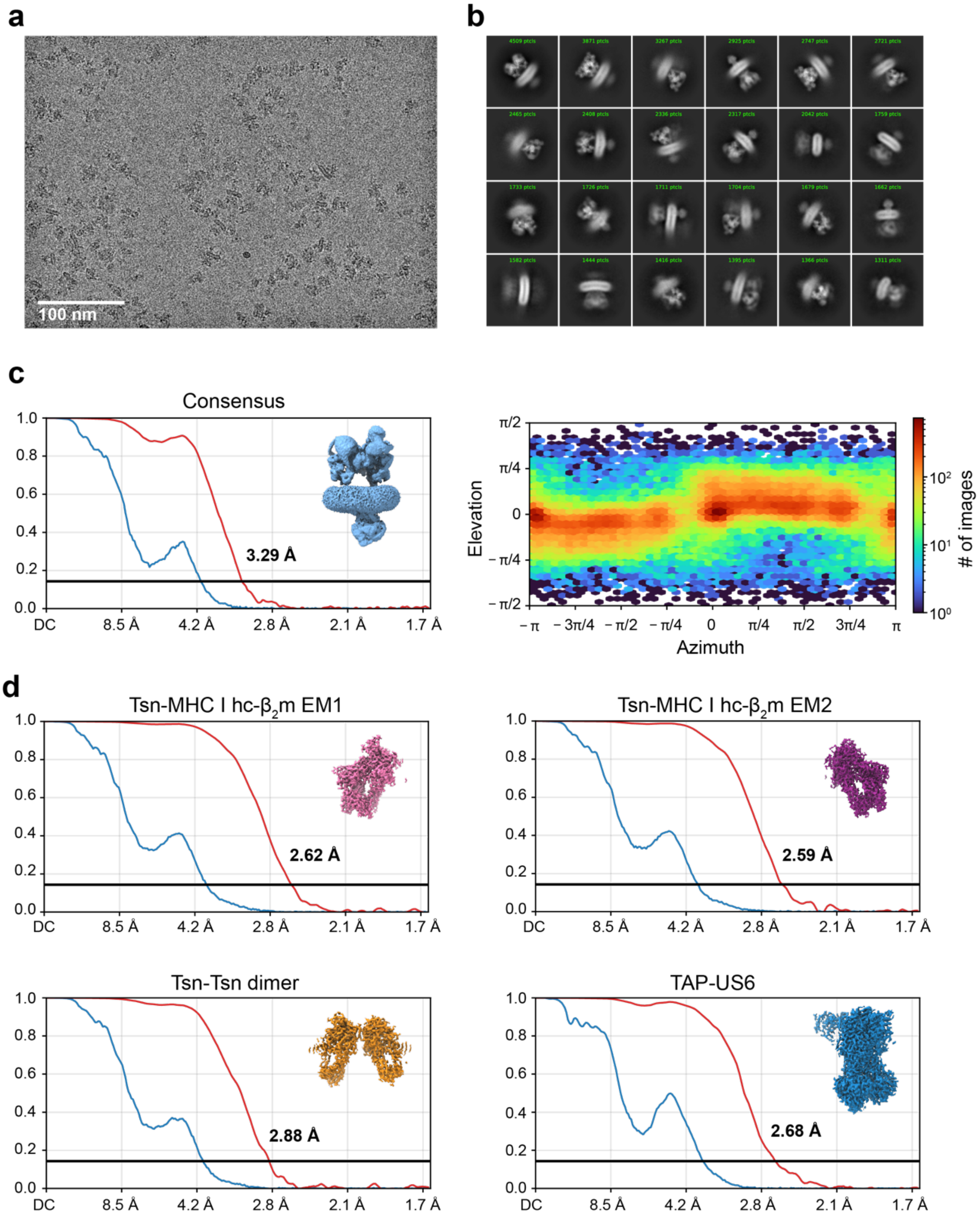


**Extended Data Fig. 3 | Single particle cryo-EM structure determination. a**, Representative cryo-EM micrograph selected from a total of 25,454 micrographs. **b**, 2D class averages of Topaz-picked particles, organized by particle population. **c**, Fourier shell correlation (FSC) curve (left) and direction distribution plot (right) of the consensus map by non-uniform refinement. The corresponding cryo-EM map is shown in the inset. **d**, FSC curves from local refinements. Corresponding focus maps are presented in the insets.

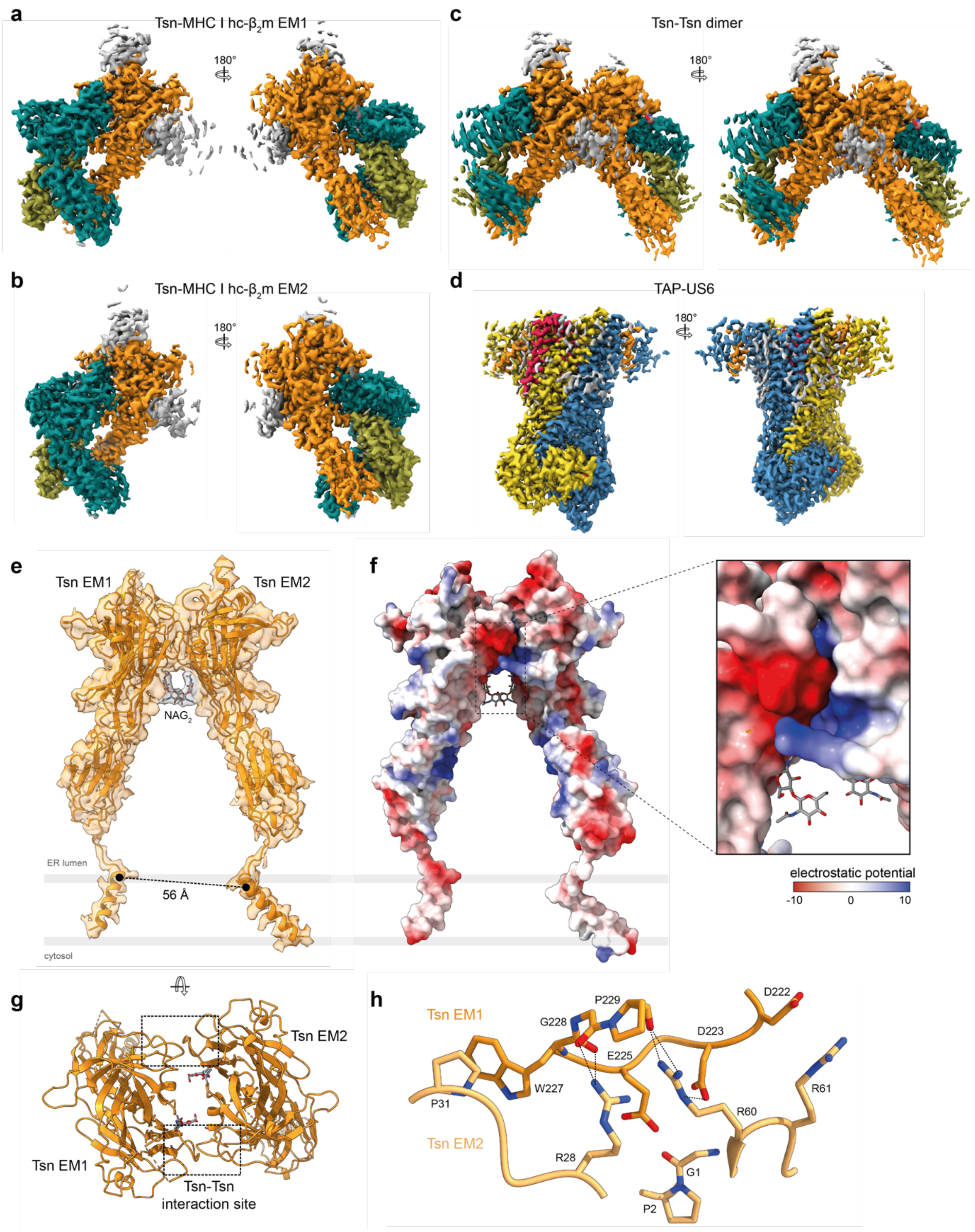


**Extended Data Fig. 4 | Focus maps used for model building, refinement, and assembly of the tapasin dimer. a-d**, Two views of the high-resolution focus maps used for model building and generation of the composite map: tapasin-MHC I heavy chain-$\beta_2$-microglobulin in editing module 1 (Tsn-MHC I hc-$\beta_2$m EM1) (**a**), the tapasin-tapasin dimer (Tsn-Tsn) (**b**), Tsn-MHC I hc-$\beta_2$m EM2 (**c**), and the antigen translocation module comprising US6, TAP1/2, and the transmembrane domain of tapasin (TAP-US6) (**d**). Individual PLC subunits are highlighted according to the color scheme in Fig. 1. **e**, Structural organization of the two editing modules via the Tsn EM1/EM2 dimer. The two Nacetylglucosamine disaccharides (NAG-NAG) of the two N233-linked N-glycans are shown as gray sticks. The distance of the transmembrane anchor points is indicated. **f**, Electrostatic charge compensation at back-to-back interaction interface (733 Å$^2$) of the two tapasin molecules (dashed box). Charged patches are shown as a Coulombic energy surface, visualizing the electrostatic potential (-10 to 10 kcal/mol). **g**, Detailed view of the Tsn-Tsn interface highlighting the two Tsn-Tsn interaction sites (dashed boxes). **h**, Close-up view on one interaction site of Tsn EM1 (orange) and Tsn EM2 (yellow orange). Interacting residues are shown as sticks and labeled; H-bond interactions are indicated with dotted lines.

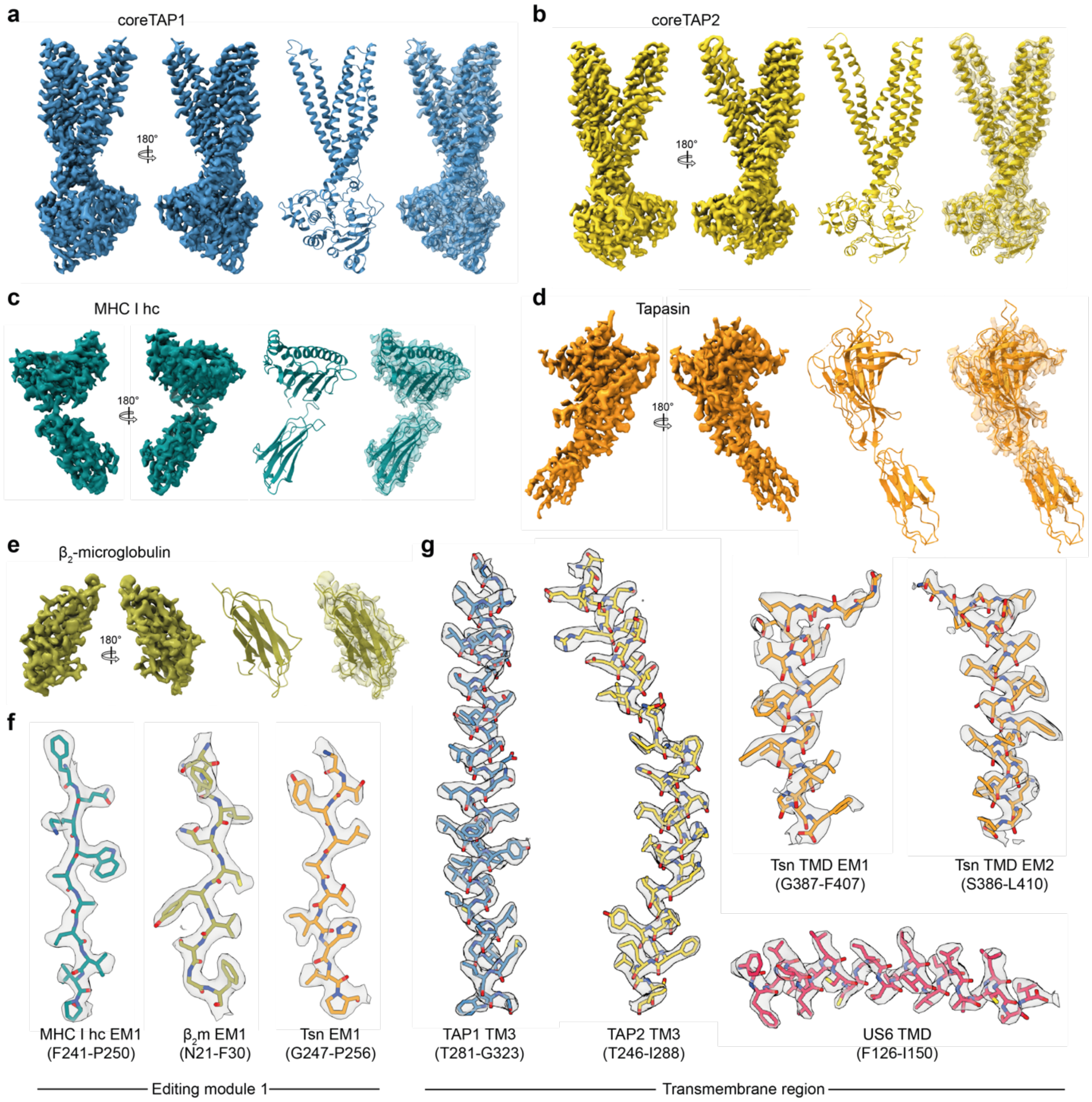


**Extended Data Fig. 5 | Map/model fit. a-e**, Single PLC subunits of the translocation module and editing module 1 highlight the quality of the map/model fit. For each subunit, two views onto the density map, the according atomic model in ribbon representation and a map-model overlay (map transparency 75%) is shown. PLC subunits are labeled and color coded as shown in Fig. 1. **f**, **g**, Cryo-EM densities (transparent gray surface) are shown with corresponding segments of the atomic models rendered in stick representation.

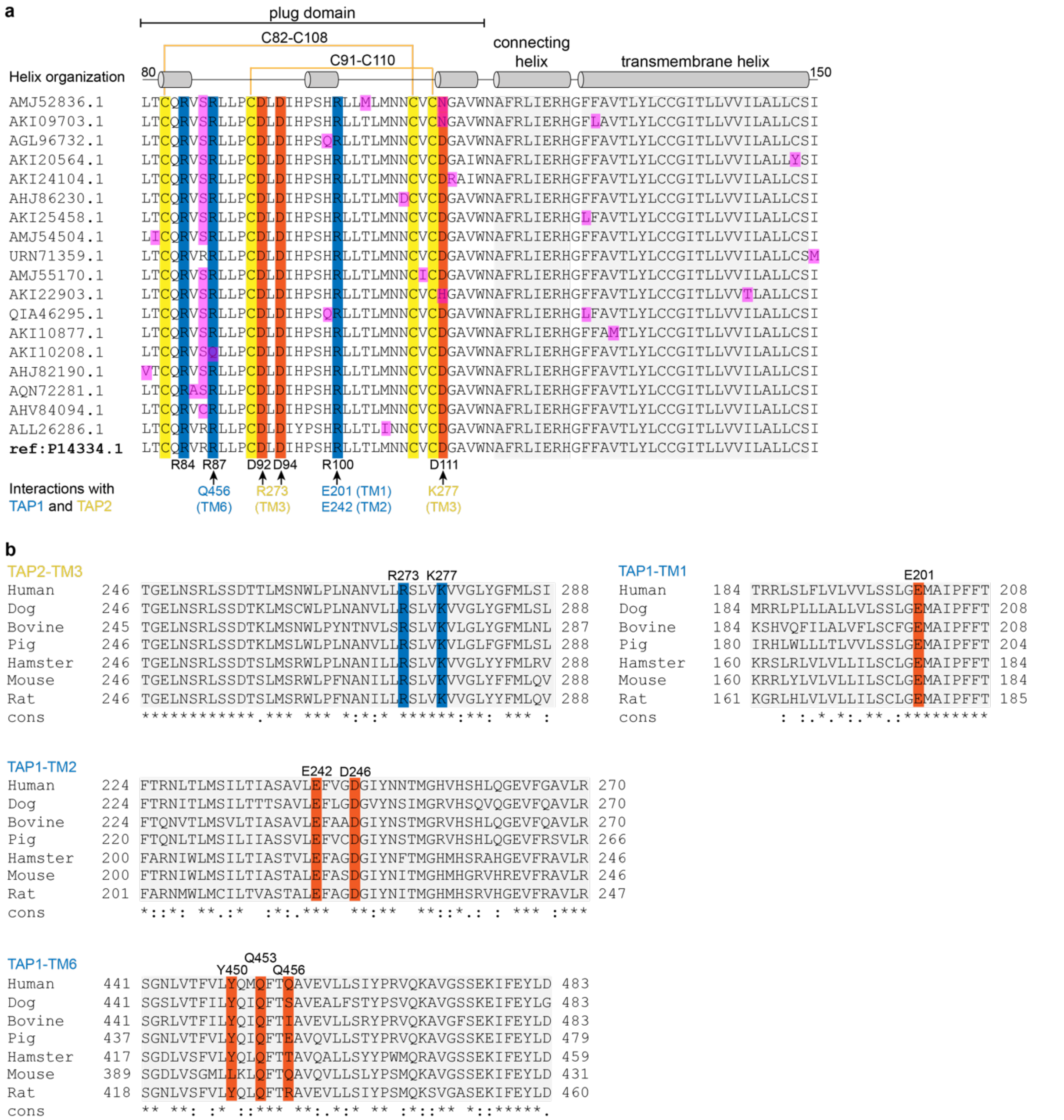


**Extended Data Fig. 6: Multiple sequence alignment of US6 from diverse human cytomegalovirus strains and of US6 interacting residues of TAP across species. a,** Sequence alignments provide insights into conserved and variable regions. Sequence variations among US6 variants are marked with magenta boxes. Amino acid sequences of the plug domain, connecting helix, and transmembrane domain of US6 are shown. Conserved cysteine residues forming two disulfide bonds (C82-C108 and C91-110) are highlighted in yellow; conserved arginine residues (R84, R87, and R100) in blue; and conserved aspartate residues (D92, D94, and D111) in red. Interactions with conserved residues in transmembrane (TM) helices of TAP1 (light blue) and TAP2 (yellow) are annotated below (see Fig. 3c,d). **b**, US6-TAP interacting residues in transmembrane helix (TM) 3 of TAP2 and TM1, TM2, and TM6 of TAP1 (Fig. 3c,d) are highlighted in blue and red boxes. cons, consensus sequence.

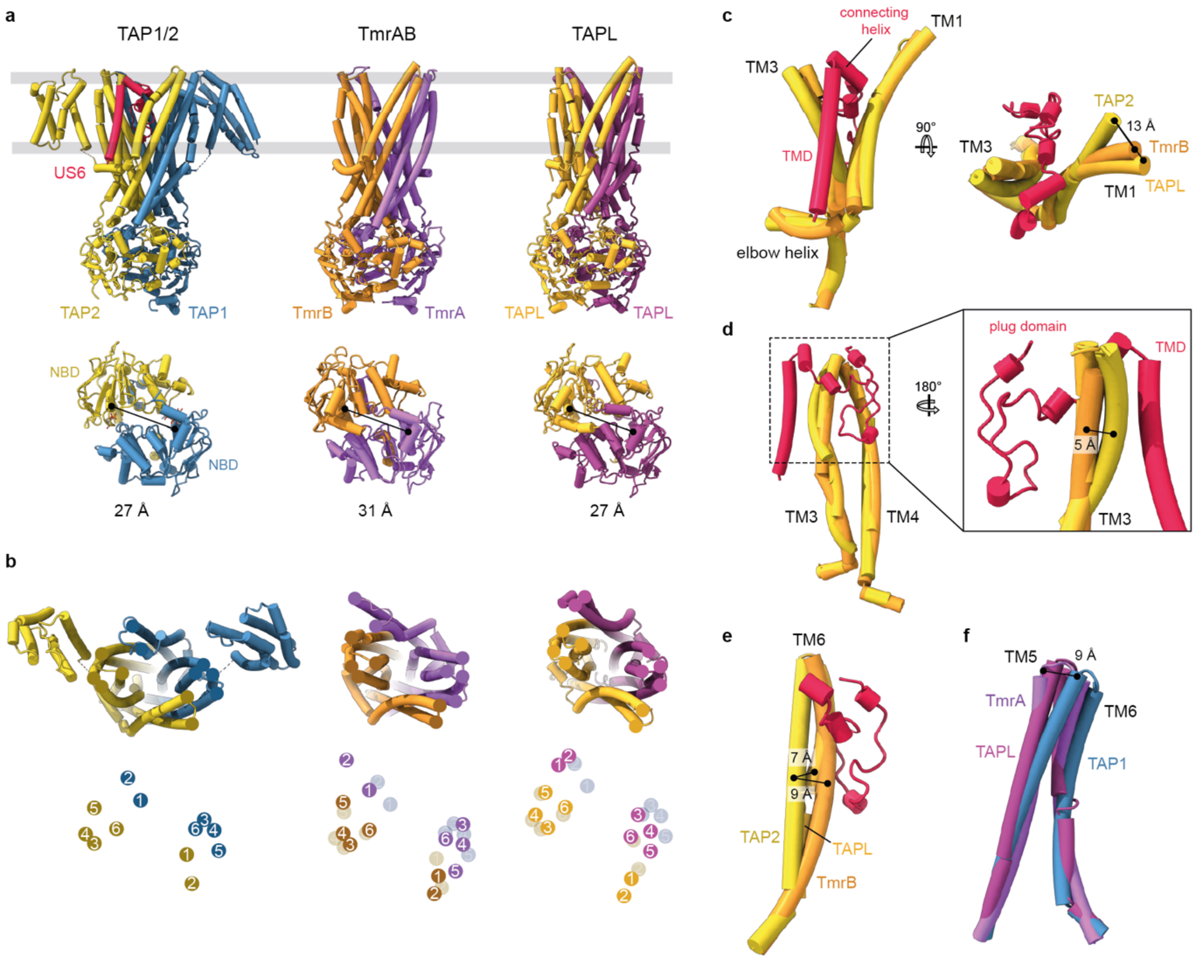


**Extended Data Fig. 7: Conformations of the US6-inhibited TAP translocation module compared to TAP-related transporters. a**, Structural comparison of the US6-bound TAP with the outward-facing open conformations of the bacterial transporter TmrAB (PDB: 6RAH)[23] and the human transporter TAPL (PDB: 7V5C)[28]. In the presence of US6, TAP adopts a semi-open conformation that is incompatible with substrate translocation. **b**, View from the ER lumen showing the positions of transmembrane (TM) helix tips, highlighting structural deviations from US6-bound conformation. **c-e**, Close-up views of altered TM helix conformations in TAP. The lumenal tip of TM1 in coreTAP2 appears displaced relative to TAP-related transporters, likely to accommodate the connecting helix of US6 (**c**). TM helices 3 and 6 of coreTAP2 display altered curvature where they interface with the US6 plug domain (**d**, **e**). **f**, The ER-lumenal ends of TM5 and 6 are bent away from the conformations observed in TmrAB and TAPL. All helices are labeled accordingly.

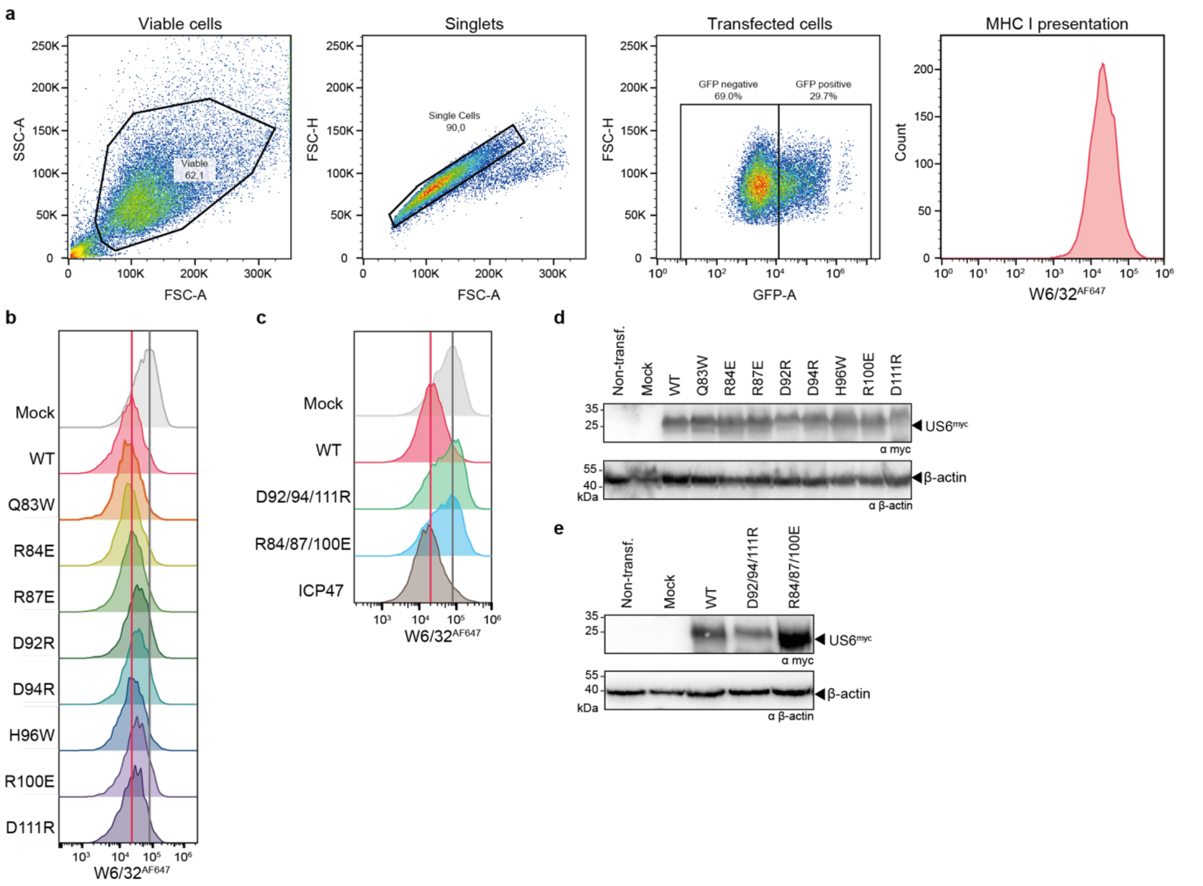


**Extended Data Fig. 8: Effect of US6 mutations on MHC I surface presentation. a**, Flow cytometry analysis of MHC I surface expression and gating strategy. HeLa cells were transfected with the plasmid pAMI_IRES_eGFP encoding myc-tagged US6 variants. Enhanced green-fluorescent protein (GFP) served as a reporter for the transfection efficiency. Gating strategy on viable cells (left), singlets (middle left), transfected, GFP-positive cells (middle right), and MHC I surface expression detected by the pan-MHC-I antibody W6/32$^{AF647}$ (right). SSC-A, side scatter - area; FSC-A/H, forward scatter – area/height; GFP-A, GFP signal - area. **b**, Cells expressing US6 wild-type or single substitutions were stained with the anti-MHC I antibody W6/32$^{AF647}$. Median fluorescence intensities (MFI) were summarized in Fig. 4f. **c**, Cells expressing US6 wild-type, triple substitutions, or the herpes simplex virus TAP inhibitor ICP47 were similarly stained with W6/32$^{AF647}$. Median fluorescence intensities (MFI) were shown in Fig. 4f. **d**, **e**, SDS-PAGE and immunoblot analysis of the transfected cells analyzed in **b** and **c**, respectively, confirming comparable expression levels of US6 variants relative to wild-type US6. β-actin was used as a loading control. Non-transf., non-transfected cells; mock, mock-transfected cells.

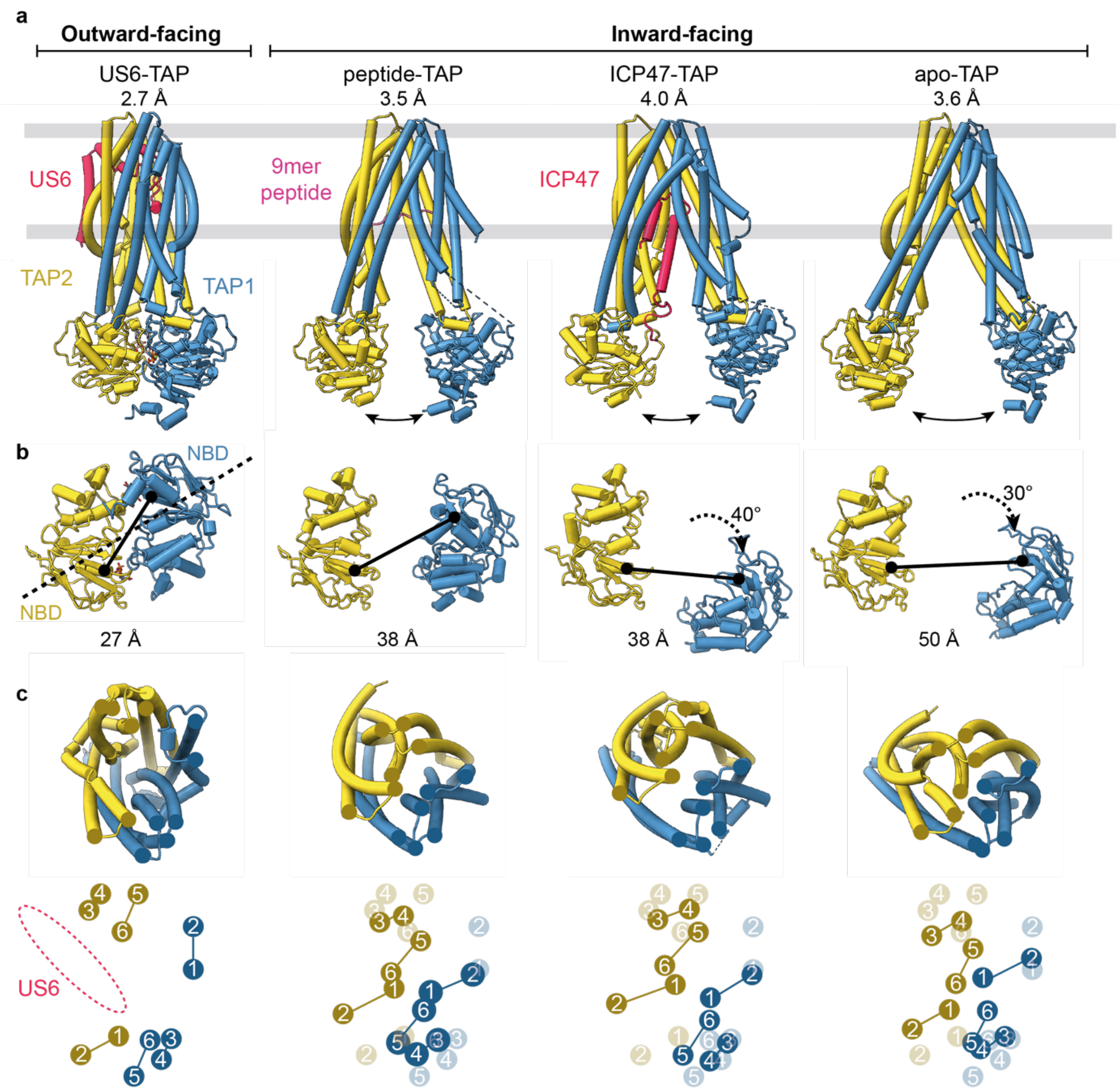


**Extended Data Fig. 9: Comparison of US6- and ICP47-inhibited TAP with apo and peptide-bound state. a**, Human cytomegalovirus US6 (red) arrests the transporter associated with antigen processing (TAP) in an outward-facing open conformation by laterally latching its transmembrane helix onto TAP2 (yellow) and plugging the lumenal exit of TAP via its central disulfide-rich domain. In contrast the herpes simplex virus ICP47 (red) inserts from the cytosolic side, wedging into the inward-facing conformation distinct from the apo and peptide-bound state of TAP (US6-PLC: PDB 9RCV; peptide-TAP[25]: 8T4F; ICP47-TAP[37]: 5U1D; apo-TAP[25]: 8T46). **b**, Superposition of TAP conformers illustrates distinct arrangements of the nucleotide-binding domains (NBDs) of apo, peptide-bound, and ICP47-inhibited TAP display different degrees of openings: while apo, peptide-bound, and ICP47-inhibited TAP show varying degrees of NBD separation, US6-TAP in the fully assembled PLC displays fully closed NBDs with asymmetrically trapped nucleotides. **c**, Conformational differences in the transmembrane (TM) helices viewed from the endoplasmic reticulum (ER) lumen, highlighting the impact of US6- versus ICP47-mediated inhibition.

**Extended Data Movie 1: Conformational dynamics of the PLC.** 3D variability analysis of PLC editing modules was performed in cryoSPARC. Three variability modes are exemplarily presented to highlight conformational dynamics within the PLC editing modules. Each series contains 20 3D structures played as oscillating frames.

**Extended Data Table 1. Cryo-EM data collection, refinement and validation statistics.**

| | PLC Consensus EMD-53326 | TAP-US6 Focus EMD-53332 | Tsn-MHC I hc-β2m EM1 Focus EMD-53334 | Tsn-MHC I hc-β2m EM2 Focus EMD-53331 | Tsn-Tsn-dimer Focus EMD-53330 |
|---|---|---|---|---|---|
| **Data collection and processing** | | | | | |
| Magnification | 105,000x | 105,000x | 105,000x | 105,000x | 105,000x |
| Voltage (kV) | 300 | 300 | 300 | 300 | 300 |
| Electron exposure (e–/Å$^2$) | 58 | 58 | 58 | 58 | 58 |
| Defocus range (μm) | -0.5 to -1.5 | -0.5 to -1.5 | -0.5 to -1.5 | -0.5 to -1.5 | -0.5 to -1.5 |
| Pixel size (Å) | 0.83 | 0.83 | 0.83 | 0.83 | 0.83 |
| Symmetry imposed | C1 | C1 | C1 | C1 | C1 |
| Initial micrographs (no.) | 25,454 | 25,454 | 25,454 | 25,454 | 25,454 |
| Final micrographs (no.) | 25,292 | 25,292 | 25,292 | 25,292 | 25,292 |
| Initial particle images (no.) | 726,723 | 726,723 | 726,723 | 726,723 | 726,723 |
| Final particle images (no.) | 109,445 | 269,213 | 269,213 | 269,213 | 269,213 |
| Map resolution (Å) | 3.29 | 2.68 | 2.62 | 2.59 | 2.88 |
| FSC threshold | 0.143 | 0.143 | 0.143 | 0.143 | 0.143 |
| Map sharpening *B* factor (Å$^2$) | -76.1 | -84.5 | -83.1 | -80.6 | -87.7 |

| | PLC Composite EMD-53326 PDB 9RCV | TAP-US6 Focus EMD-53332 PDB 9RCV | Tsn-MHC I hc-β2m EM1 Focus EMD-53334 PDB 9RCV | Tsn-MHC I hc-β2m EM2 Focus EMD-53331 PDB 9RCV | Tsn-Tsn-dimer Focus EMD-53330 PDB 9RCV |
|---|---|---|---|---|---|
| **Refinement** | | | | | |
| Model composition | | | | | |
| Non-hydrogen atoms | 23535 | 11314 | 5906 | 5981 | 5707 |
| Protein residues | 2993 | 1504 | 740 | 751 | 745 |
| ATP/ADP | 2 | 2 | 0 | 0 | 0 |
| Carbohydrates | 4 | 0 | 2 | 2 | 4 |
| Waters | 4 | 4 | 0 | 0 | 0 |
| Ions | 2 | 2 | 0 | 0 | 0 |
| *B* factors (Å$^2$) | | | | | |
| Protein | 67.65 | 69.98 | 64.04 | 65.84 | 63.68 |
| Ligand | 70.16 | 64.77 | 94.04 | 73.04 | 68.31 |
| R.m.s. deviations | | | | | |
| Bond lengths (Å) | 0.004 | 0.004 | 0.004 | 0.004 | 0.004 |
| Bond angles (°) | 0.967 | 0.961 | 0.975 | 0.980 | 1.030 |
| **Validation** | | | | | |
| MolProbity score | 0.82 | 0.82 | 0.75 | 0.86 | 0.96 |
| Clashscore | 1.11 | 1.14 | 0.78 | 1.28 | 1.32 |
| Poor rotamers (%) | 0.64 | 0.99 | 0.16 | 0.48 | 0.17 |
| Ramachandran plot | | | | | |
| Favored (%) | 98.58 | 98.89 | 98.48 | 97.98 | 97.52 |
| Allowed (%) | 1.42 | 1.11 | 1.52 | 2.02 | 2.48 |
| Disallowed (%) | 0.00 | 0.00 | 0.00 | 0.00 | 0.00 |
